# Large-scale digital traces of university students show that morning classes are bad for attendance, sleep, and academic performance

**DOI:** 10.1101/2021.05.14.444124

**Authors:** Sing Chen Yeo, Clin K.Y. Lai, Jacinda Tan, Samantha Lim, Yuvan Chandramoghan, Joshua J. Gooley

## Abstract

Attending classes and sleeping well are important for students’ academic success. However, early classes might impede learning by contributing to absenteeism and insufficient sleep. We used big datasets collected passively from university students to test the hypothesis that morning classes are associated with poorer attendance, shorter sleep, and lower grades. Wi-Fi connection data were used to estimate attendance rates of 24,678 students enrolled in lecture courses with start times ranging from 08:00 to 16:00. Students’ interactions with the university’s Learning Management System (LMS) were used to estimate nocturnal sleep opportunities by compiling 17.4 million logins from 39,458 students with data sorted by students’ first class of the day. Objective sleep behavior was assessed in 181 students who took part in a 6-week actigraphy study. We found that Wi-Fi confirmed attendance was about 15 percentage points lower in students taking classes at 08:00 compared with later start times. Actigraphy data revealed that students frequently slept past the start of morning classes. LMS and actigraphy data showed that nocturnal sleep opportunities and total sleep time decreased with earlier class start times due to students waking up earlier. Analyses of grades in 27,281 students showed that having morning classes on more days of the week resulted in a lower grade point average. These findings suggest cumulative negative effects of morning classes on learning. Early morning classes force many students to decide to either sleep more and skip class, or sleep less to attend class. Therefore, universities should avoid scheduling early morning classes.

**Significance Statement:** We show that morning classes are associated with lower attendance, shorter nocturnal sleep, and lower grade point average in university students. Scalable methods for measuring attendance and sleep were developed using students’ Wi-Fi connection data and interactions with the Learning Management System. Students had lower attendance rates and frequently slept past the start of early morning classes. However, students still lost about an hour of sleep on average when they had early morning classes due to waking up earlier than usual. Students who had morning classes on more days of the week had a lower grade point average. Our results suggest cumulative negative effects of morning classes on students’ academic performance. Universities should avoid scheduling early morning classes.

## Introduction

University students who regularly attend classes and sleep well are more likely to get good grades (1-4). Attending classes increases students’ interactions with instructors and classmates, and provides structured time for covering key learning points. Unsurprisingly, class attendance is among the strongest predictors of academic achievement (1). Sleeping well is also important for optimizing cognitive performance and readiness to learn. Inadequate sleep impairs attention and memory processes (3, 5-7), which may prevent students from reaching their full learning potential in class (i.e., presenteeism). Moreover, feeling tired and oversleeping are frequently cited as reasons why university students skip classes (8-11). Effects of absenteeism and presenteeism on grades may have long-term consequences on students’ employment opportunities (12), job performance ratings (13), and salary (14). Therefore, universities should adopt practices that improve students’ attendance rates and sleep behavior to position them to succeed in the classroom and workforce.

Growing evidence indicates that early class start times are detrimental for students’ sleep and daytime functioning. During adolescence and early adulthood, environmental and biological factors result in a delay in the preferred timing of sleep (15, 16). Hence, students who go to bed late and must wake up early for class have shorter nocturnal sleep (17). The circadian drive for sleep may also reach its peak close to the time that students are expected to attend early morning classes. The combined effects of short sleep and circadian misalignment can lead to daytime sleepiness and impaired cognitive performance (18). Delaying the start time of high schools has been shown to increase sleep duration and decrease sleepiness by allowing adolescents to sleep in longer (19-23). However, there are mixed findings regarding the benefits of starting school later on absenteeism and academic outcomes (20, 22, 24). It is also unclear if results for high schools are generalizable to universities.

It is important to study effects of class start times on behavior in university students because they face environmental pressures that are different compared with high school. Many university students are living away from home for the first time and encounter new social contexts, demanding coursework, and opportunities for late-night socializing (25). The increased autonomy in how university students spend their time may lead to later bedtimes on school nights compared to when they were in high school (26). University students also have a less structured timetable in which the timing of their first class of the day can vary across the school week. This could lead to daily changes in students’ wake-up times and nocturnal sleep duration (27). Class attendance is also rarely monitored for lectures or seminars at universities, whereas attendance is compulsory and tracked in high schools. University students therefore have the freedom to skip classes and may decide, for example, to sleep instead of going to early morning classes. This, in turn, could impact students’ grades (28-32).

Universities need scalable methods for evaluating the impact of class start times on students’ attendance and sleep behavior. Class scheduling practices are unlikely to change without university-wide evidence of a problem. Most studies on attendance and sleep have been limited to convenience samples with small numbers of courses or students. Recent work suggests that it may be possible to monitor students’ class attendance by detecting whether they connect to a Wi-Fi access point in the classroom (33, 34). Given that nearly all students carry a Wi-Fi enabled device (e.g., smartphone, laptop, or tablet), Wi-Fi connection data could be used across the entire university to estimate class attendance without the need for active participation by students or instructors. Students’ digital traces on university learning platforms could also be used to estimate their diurnal sleep behavior. Many universities use a Learning Management System (LMS) as the primary online platform for students to download course materials, submit assignments, complete quizzes, and participate in class discussions. Students’ interactions with the LMS represent a form of wake signal that could be used to determine when sleep is likely to occur (30). If LMS data can be shown to provide a reliable estimate of sleep timing, universities can use this information for measuring the aggregate impact of class start times on sleep behavior of the entire student body.

The objective of our study was to develop scalable methods for testing the effects of class start times on students’ attendance and sleep behavior. At a large university, Wi-Fi connection data were used to estimate students’ lecture attendance rates, and LMS data were used to estimate sleep opportunities. In parallel, an actigraphy study was conducted to validate and extend the findings. First, we tested the hypothesis that early morning classes are associated with lower Wi-Fi confirmed attendance rates. Actigraphy data were used to determine whether students chose to sleep instead of attending early classes. Second, we tested the hypothesis that early morning classes are associated with earlier wake-up times and shorter sleep. This was assessed using LMS and actigraphy data sorted by students’ first class of the day. Third, we tested the hypothesis that morning classes are associated with lower course grades, and that students who have morning classes more days of the week have a lower cumulative grade point average.

## Results

### Class start times and students’ attendance

Students’ class attendance rates were estimated using time and location data from their Wi-Fi connection logs (**Table S1; Fig. S1**). Students were confirmed as present if they connected to a wireless access point in the classroom during class hours. First, we showed that there was a strong linear correlation between instructor-reported attendance and Wi-Fi confirmed attendance (53 class sessions; Pearson’s r = 0.98), indicating that Wi-Fi connection data can be used as a reliable indicator of class attendance (**Fig. 1A**). Next, this method was used to measure Wi-Fi confirmed attendance rates for 24,678 unique students enrolled across 337 large lecture courses (≥100 students enrolled per course) with class start times ranging from 08:00 to 16:00 (**Fig. 1**; **Fig. S1**). There was a significant association between Wi-Fi confirmed attendance and class start time (ANOVA, *F*_5,36807_ = 239.9, *P* < 0.001). The attendance rate for students with lecture classes at 08:00 was about 15 percentage points lower compared with students whose lecture classes started at 10:00 or later (**Fig. 1B**). In within-student comparisons of Wi-Fi confirmed attendance rates, there was a medium effect size of starting class at 08:00 relative to all other class start times (Cohen’s d, range = 0.45 to 0.53; *P* < 0.001 for all pairwise comparisons) (**Fig. 1C**). There was little to no effect of class start time on attendance for all other pairwise comparisons (Cohen’s d, range = -0.14 to 0.16).

**Figure 1.**
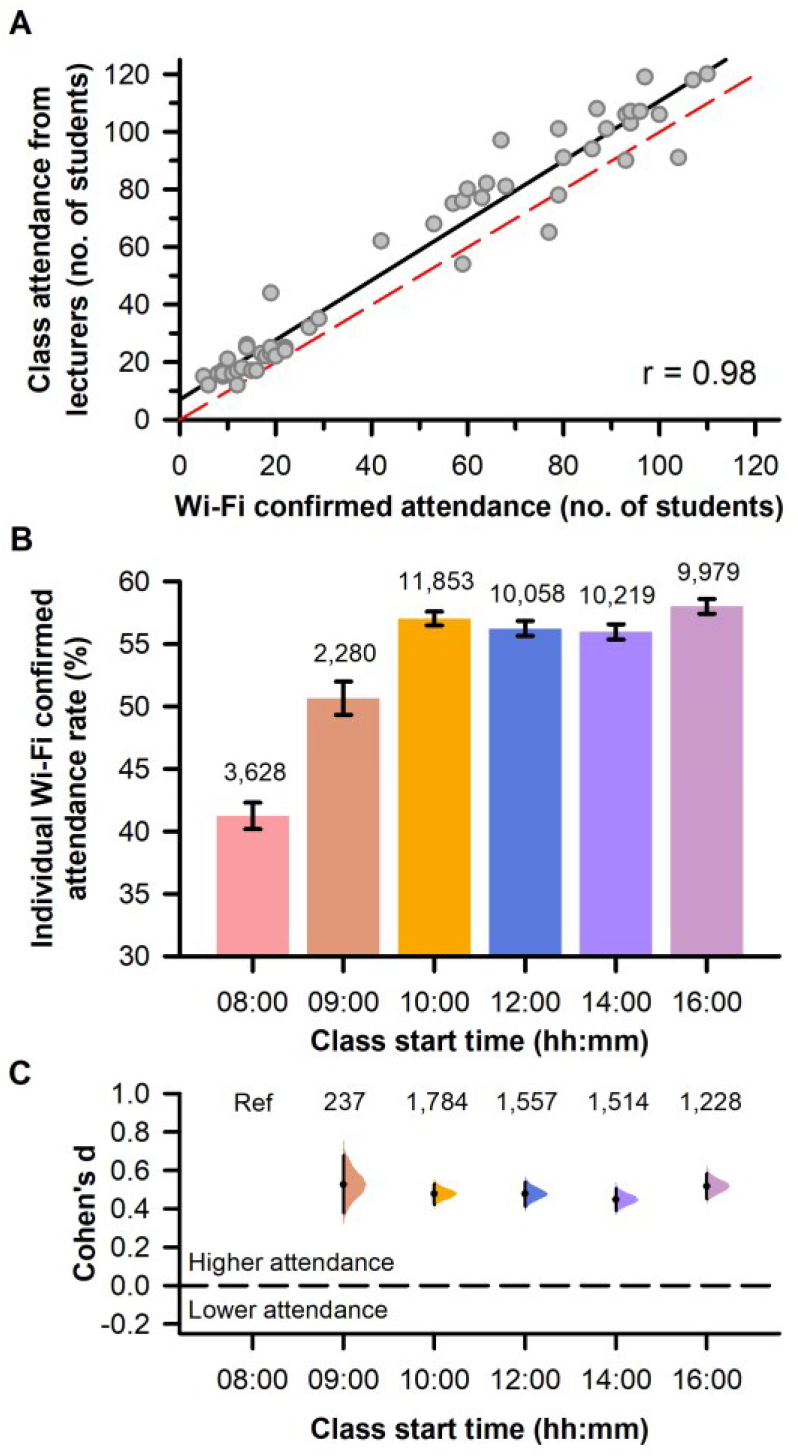
Wi-Fi confirmed lecture attendance was lower for early morning classes. (**A**) Instructor-reported attendance was strongly correlated with Wi-Fi confirmed attendance. Each circle shows attendance data for an individual class session (53 class sessions across 13 different courses). The black trace shows the best-fit linear regression line and the red dashed trace is the unity line. (**B**) Individually-determined Wi-Fi confirmed attendance rates are shown for different class start times. Sample sizes are displayed at the top of each bar for each start time. The mean and 95% CIs are shown. (**C**) Effect sizes (Cohen’s d) are shown for within-student comparisons of Wi-Fi confirmed attendance rates. Effect sizes were determined for each class start time relative to 08:00. The number of students in each comparison is indicated at the top of the plot. The paired mean difference for each comparison is shown with 95% CIs and the bootstrap sampling distribution.

Next, we examined whether the lower attendance rate for early classes was driven by students sleeping instead of going to class. Sleep behavior was assessed in 181 students who took part in a 6-week actigraphy study during the school semester (**Fig. 2A**). Data for 6,546 sleep offset times were sorted by students’ first class of the day for individuals who provided information on their daily commute time to school (n = 174) (**Fig. 2B**). We then assessed the frequency of instances whereby students woke up after the start of their first class, or woke up before class but could not have reached on time based on their self-reported travel time. The frequencies of sleeping past the start of class (**Fig. 2C**) and waking up too late to reach class on time (**Fig. 2D**) increased with earlier start times (*χ*^2^ = 394.4, *P* < 0.001 and *χ*^2^ = 487.5, *P* < 0.001, respectively). Students did not wake up in time for nearly one third of classes that took place at 08:00, whereas they rarely slept past the start of classes that began at noon or later. Using Wi-Fi connection data, we explored whether students who woke up late were detected on campus during their class. Students who slept past the start of their 08:00 class, or woke up too late to reach on time, rarely connected to the campus Wi-Fi network (11.2% and 18.4% of all instances, respectively), suggesting that they did not attend class. In contrast, students who woke up in time to reach their 08:00 class were usually detected (69.6% of instances; *χ*^2^ = 322.0, *P* < 0.001).

**Figure 2.**
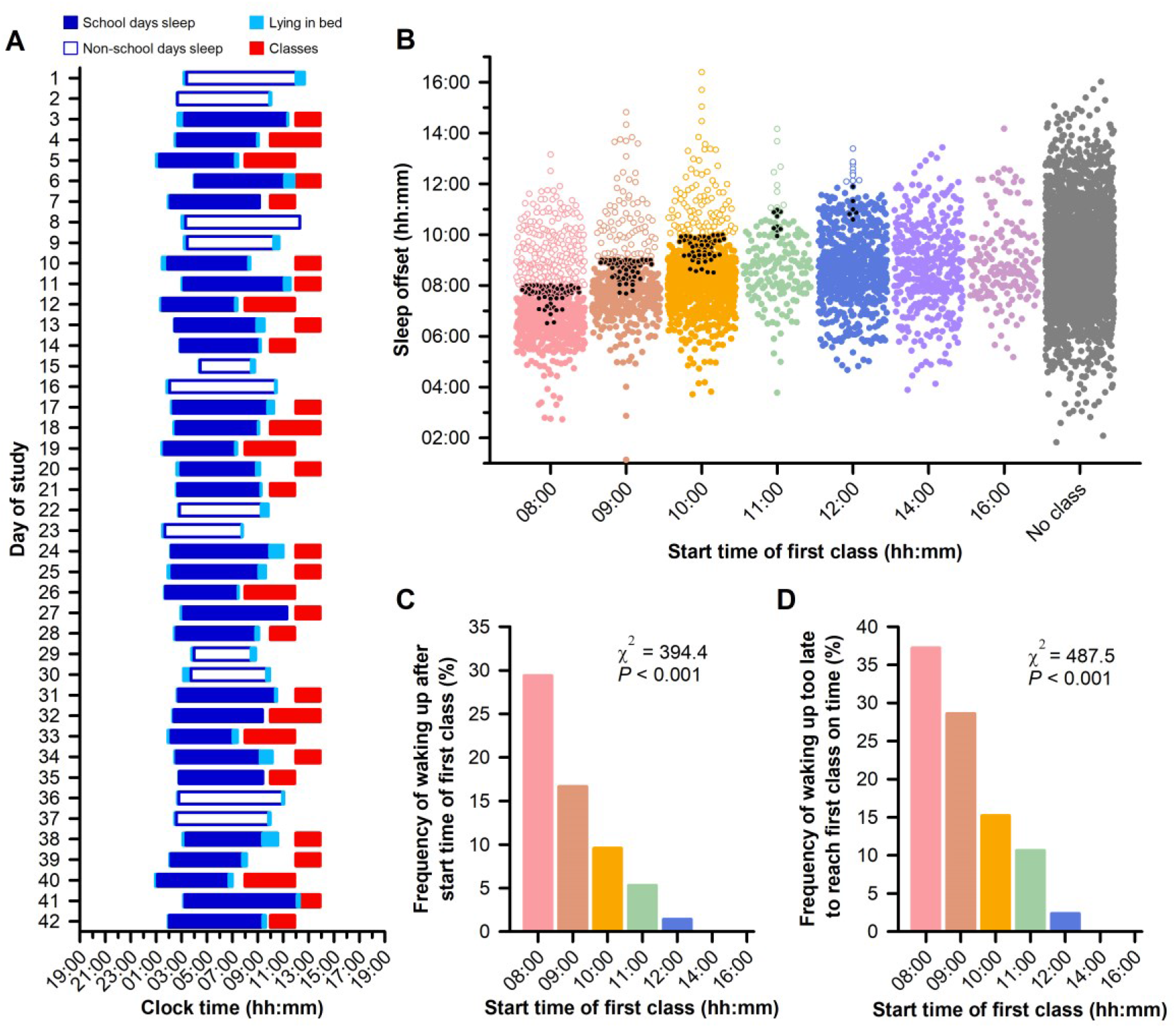
Students frequently slept past the start of morning classes. (**A**) Sleep periods and scheduled classes are shown in a representative student who took part in a 6-week actigraphy study. (**B**) Sleep offsets from 174 students are sorted by students’ first class of the day. Each circle corresponds to an individually-determined sleep offset value. Open circles show instances when students woke up after the start of their class. Black circles show instances when students did not wake up early enough to reach class on time when their self-reported travel time was taken into account. The frequencies of (**C**) waking up after the start of class, and (**D**) waking up too late to reach class on time are shown for different class start times.

### Class start times and students’ sleep opportunities

Students’ opportunities for nocturnal sleep were investigated for different class start times by aggregating their LMS login data over 5 semesters (**Table S1**). Given that students must be awake to log in to the LMS, we used this ‘wake signal’ to assess the impact of class start times on the time window when nocturnal sleep could occur. Diurnal time courses of LMS logins were constructed by compiling 17.4 million time-stamped logins from 39,458 students (**Fig. 3A**; **Fig. S2**). LMS login data on school days were sorted by students’ first class of the day and compared with data on non-school days (**Fig. 3A**). The diurnal pattern of login activity on school days comprised a 24-hour component with higher activity during the daytime and hourly peaks that corresponded to the timing of classes. The time courses of LMS logins for different class start times were highly reproducible across semesters (**Fig. S2**). On school days with earlier start times (08:00, 09:00 and 10:00), LMS login activity increased earlier in the morning than days with no classes. By comparison, the 24-h diurnal component of login behavior was similar between school days and non-school days when students’ first class of the day took place in the afternoon.

**Figure 3.**
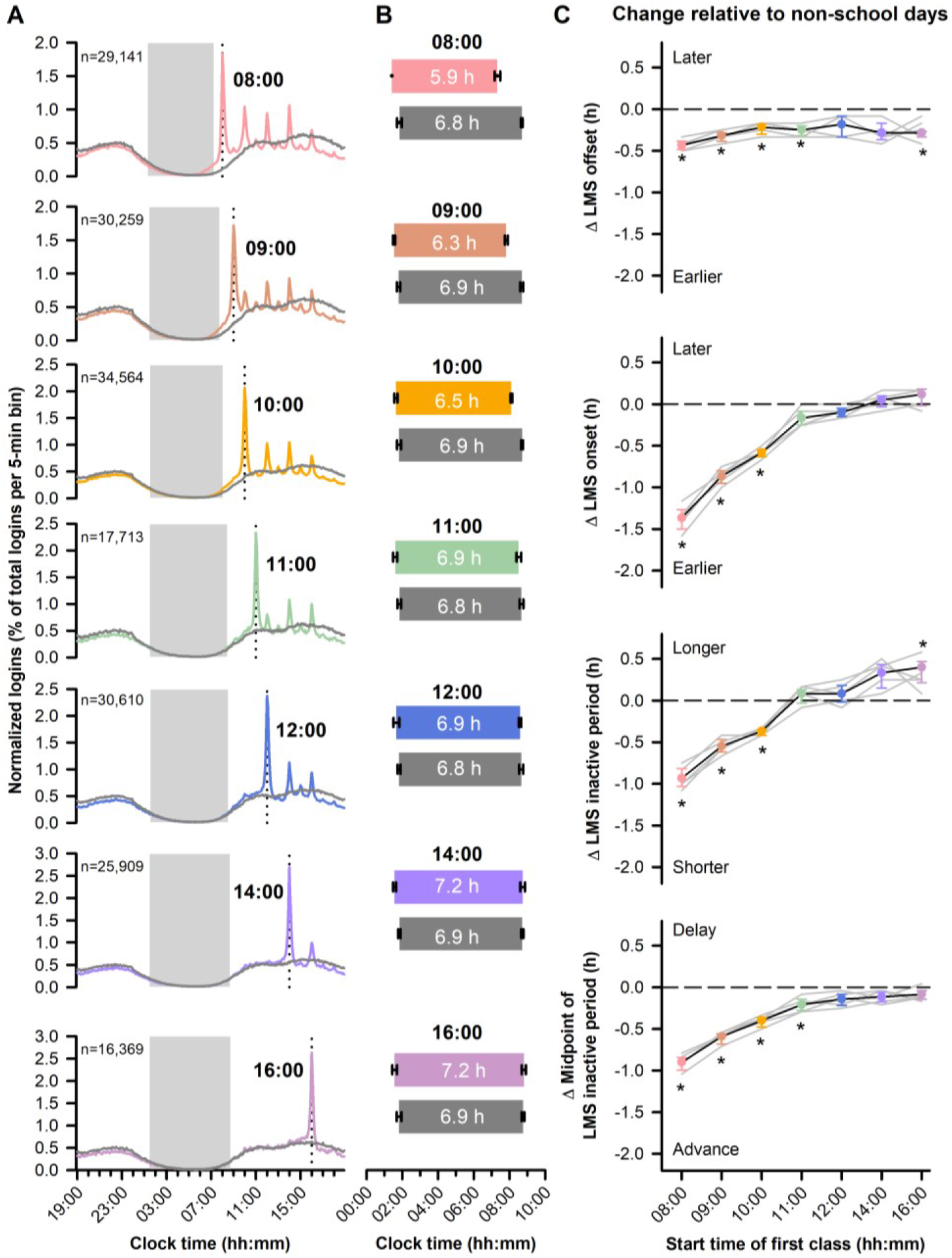
Earlier class start times resulted in shorter sleep opportunities. (**A**) Diurnal time courses of Learning Management System (LMS) logins are shown in students whose data are sorted by their first class of the day. Colored traces show data for school days and grey traces show data for non-school days in the same group of participants. The vertical dotted line in each plot shows the start time of students’ first class. The grey boxes show LMS inactive periods on school nights when login activity fell below threshold (see Methods). (**B**) LMS login onset and offset values are shown for different class start times (colored bars) and compared with data in the same students for non-school days (grey bars). The duration of the LMS inactive period is indicated in each bar. The mean ± 95% CIs are shown for data across 5 semesters. (**C**) Changes in LMS login behavior are shown for different class start times relative to non-school days. The paired mean differences are shown with 95% CIs. Grey traces show results for individual semesters. Asterisks show pairwise comparisons that reached statistical significance (*P* < 0.007, Bonferroni-adjusted).

For each semester and class start time, the offset and onset of the aggregated LMS login rhythm were determined using a threshold crossing method (See Materials and Methods). The duration and midpoint of the LMS inactive period were measured between LMS offset and onset, which spanned the nighttime. Values for each LMS-derived parameter were similar across individual semesters (**Table S2**). The LMS inactive period was shorter on nights that preceded morning classes (08:00, 09:00 and 10:00) compared with later classes or non-school days (**Fig. 3A-B**). The shorter opportunity for sleep for early morning classes was explained by an earlier LMS login onset (**Fig. 3B**).

Next, we quantified associations of class start time (08:00, 09:00, 10:00, 11:00, 12:00, 14:00, 16:00, and non-class days) with LMS login behavior across semesters. Significant associations were observed for class start time with the LMS login offset (*F*_7,28_ = 14.8, *P* < 0.001), the LMS login onset (*F*_7,28_ = 205.3, *P* < 0.001), the duration of the LMS inactive period (*F*_7,28_ = 94.4, *P* < 0.001), and the midpoint of the LMS inactive period (*F*_7,28_ = 147.2, *P* < 0.001). Relative to days with no classes, the LMS login offset occurred slightly earlier but did not vary much for different class start times (**Fig. 3C**). In contrast, the LMS login onset tracked closely the start time of morning classes, with activity starting more than an hour earlier for classes at 08:00 relative to non-school days (range = 1.17 h to 1.58 h across semesters) (**Fig. 3C**). By comparison, there was no difference in LMS login onset for class start times in the afternoon relative to non-school days. Consequently, a decrease in the LMS inactive period was observed only before morning classes. Relative to non-school days, the LMS inactive period was about an hour shorter for classes that started at 08:00 (range = -0.75 h to -1.08 h across semesters) (**Fig. 3C**). Similarly, the midpoint of the LMS inactive period occurred earlier only for morning classes, with an advance of nearly an hour on days when students had a class at 08:00 (range = 0.79 h to 1.04 h across semesters) (**Fig. 3C**).

We validated findings for LMS login behavior with actigraphy data collected from 181 undergraduates with a total of 7,312 nocturnal sleep recordings (**Fig. 4**). The diurnal time courses of activity counts (i.e. wrist movements) closely resembled findings for the LMS login rhythm (**Fig. 4A**). Earlier class start times were associated with earlier sleep offsets and decreased nocturnal total sleep time (**Fig. 4B**; **Table S3**). Significant associations were observed for class start time with sleep onset (*F*_7,548_ = 4.5, *P* < 0.001), sleep offset (*F*_7,567_ = 49.6, *P* < 0.001), nocturnal total sleep time (*F*_7,577_ = 32.4, *P* < 0.001), nocturnal time in bed for sleep (*F*_7,573_ = 33.7, *P* < 0.001), and midpoint of sleep (*F*_7,551_ = 33.2, *P* < 0.001).

**Figure 4.**
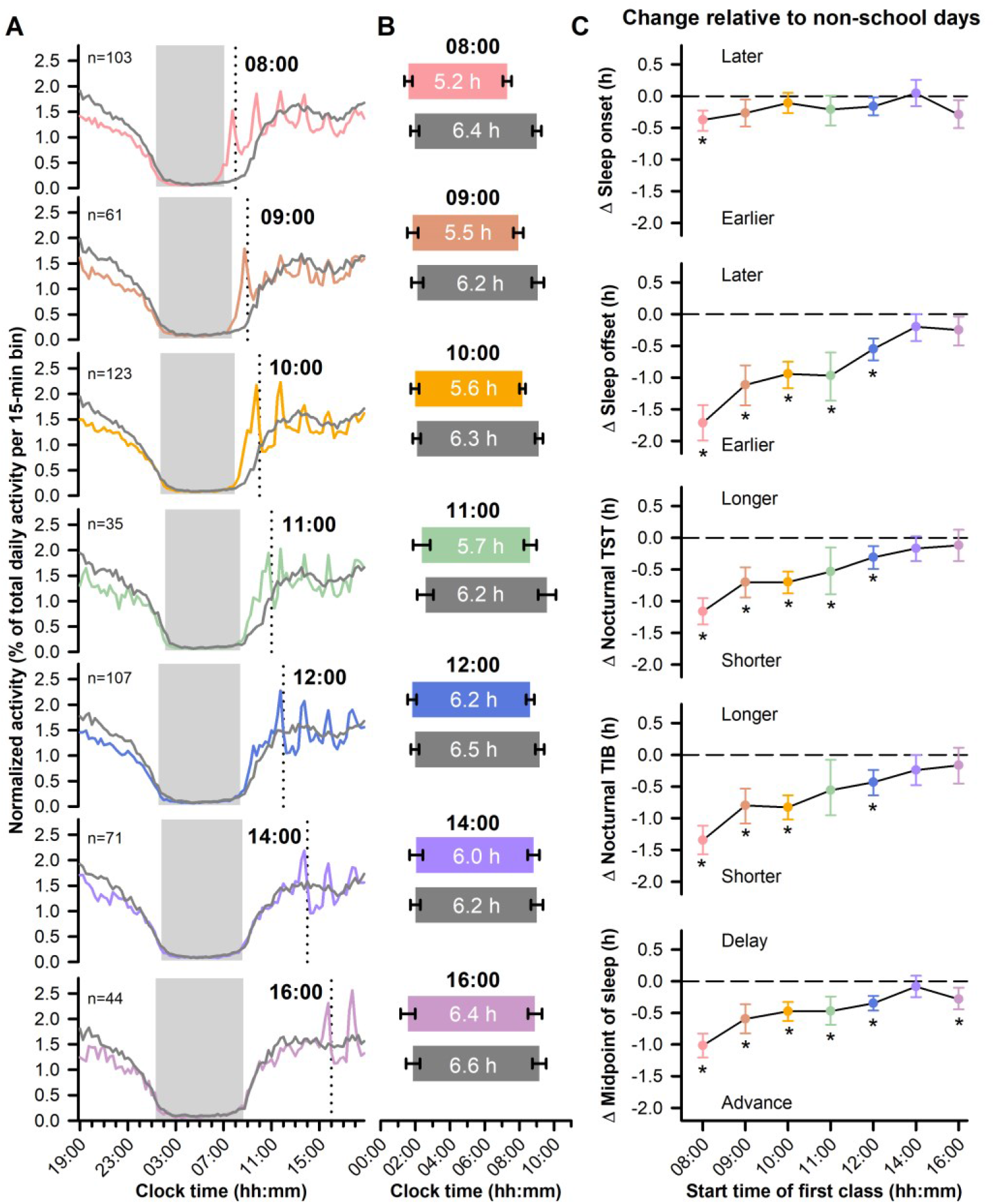
Earlier class start times were associated with shorter sleep. (**A**) The averaged diurnal time courses of actigraphy-determined activity counts are shown in students whose data are sorted by their first class of the day. Colored traces show data for school days and grey traces show data for non-school days in the same group of participants. The vertical dotted line in each plot shows the start time of students’ first class. The grey boxes show the average nocturnal sleep periods. (**B**) Sleep onset and sleep offset values are shown for different school start times (colored bars) and compared with data in the same students for non-school days (grey bars). The average nocturnal total sleep time is indicated in each bar. The mean ± 95% CIs are shown. (**C**) Changes in sleep behavior are shown for different class start times relative to non-school days. The paired mean differences are shown with 95% CIs. Asterisks show pairwise comparisons that reached statistical significance (*P* < 0.007, Bonferroni-adjusted). TIB, time in bed for sleep; TST, total sleep time.

Relative to non-school days, sleep onset occurred slightly earlier for classes that started at 08:00 but there was little variation across class start times (**Fig. 4C**). In contrast, sleep offset tracked the timing of students’ first class of the day, with larger advances for earlier morning classes (Cohen’s d, range = -1.29 to -0.75 for classes starting from 08:00 to 11:00; **Table S4**). Students whose first class took place at 08:00 advanced their sleep offset by about 1.7 h relative to non-school days (**Fig. 4C**). Both nocturnal total sleep time and time in bed for sleep decreased with earlier class start times (Cohen’s d, range = -1.14 to -0.48 for classes starting from 08:00 to 11:00; **Table S4**), and the reduction in sleep exceeded an hour in students with classes at 08:00 (**Fig. 4C**). Additionally, early class start times were associated with a greater advance in the midpoint of sleep (i.e., greater social jet lag) relative to non-school days (Cohen’s d, range = -0.89 to -0.40 for classes starting from 08:00 to 11:00; **Table S4**). Students’ midpoint of sleep occurred about an hour earlier when they had a class at 08:00 on the following day, as compared with days with no classes (**Fig. 4C**).

### Class start times and grades

The relationship between class start time and grades was analyzed in 27,281 students taking the same number of course credits (i.e., equivalent workload expressed in time units) (**Table S1**). Due to heterogeneity in the timing of classes within courses (e.g., a lecture and tutorial scheduled at different times for the same course), we categorized courses as occurring exclusively in the morning or exclusively in the afternoon (See Materials and Methods). The distribution of grades in morning courses did not differ relative to afternoon courses (Mann-Whitney U Test, U = 33, *P* = 0.218), and there was no effect of taking courses in the morning on grades relative to the afternoon (Cliff’s delta = 0.013, 95% CI = -0.003 to 0.030) (**Fig 5A; Fig S3**).

**Figure 5.**
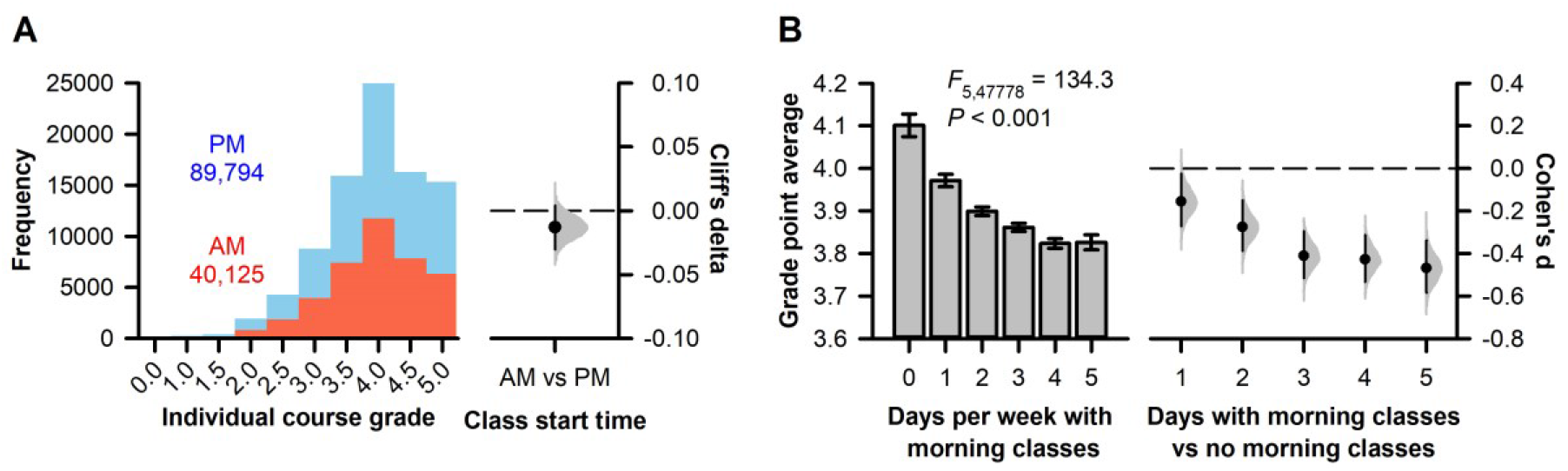
Students with morning classes on more days of the week had a lower grade point average. (**A**) The distributions of grades are shown for courses that took place exclusively in the morning (starting before 12:00) or exclusively in the afternoon (starting at 12:00 or later). The number of morning and afternoon course grades is indicated on the plot. The effect size (Cliff’s delta) of morning courses on grades is shown relative to afternoon courses. (**B**) Grade point average was lower in students who had morning classes on more days of the week. The mean ± 95% CIs are shown for students categorized by the number of days that they had morning classes. Effect sizes (Cohen’s d) with 95% CIs and the bootstrap sampling distributions are plotted for days with morning classes (1 to 5 days) versus having no morning classes.

Next, we tested whether students with more days of morning classes had a lower grade point average (e.g., due to cumulative effects of lower attendance or short sleep on overall performance). There was a significant negative association between grade point average and days of the week with morning classes (*F*_5,47778_ = 134.3, *P* < 0.001). Students with no morning classes had a higher grade point average than all other groups, despite having a comparable school workload based on course credits (**Fig. 5B**). Relative to students with no morning classes, having morning classes on 1 or 2 days of the week was associated with a small effect on grade point average (Cohen’s d = -0.16 and -0.28, respectively), and having morning classes on 3 to 5 days of the week was associated with a medium effect on grade point average (Cohen’s d = - 0.41, -0.43, and -0.47, respectively) (**Fig. 5B**). Similar findings were obtained when analyses were performed separately in each semester, demonstrating that effects of morning classes on grade point average were reproducible across semesters and academic years (**Fig. S3**).

## Discussion

Our study showed that morning classes are associated with lower lecture attendance, shorter nocturnal sleep, and lower grade point average. Many students are forced to make one of two undesirable choices when faced with early class start times: sleep more instead of attending class, or sleep less in order to attend class. Classes that started at 08:00 were especially harmful for attendance and sleep. Wi-Fi confirmed attendance rates were about 15 percentage points lower in students taking classes at 08:00 compared with later class start times. Even though students frequently slept past the start of classes at 08:00, they still lost about an hour of sleep on average. This was shown using LMS-derived sleep opportunities and confirmed by actigraphy. Students who had more days of morning classes in a week also had a lower grade point average. Our findings suggest cumulative negative effects of morning classes on absenteeism and presenteeism that lead to poorer academic achievement.

We used Wi-Fi connection data to show that students’ attendance was lowest for lecture classes in the early morning (i.e., 08:00). Our results are consistent with prior studies that compared instructor-reported attendance for the same course offered at different times of day (35-38). In large lectures/seminars and small classes, attendance was lower when classes were held in the early morning compared with later in the day. An important limitation of these studies is that attendance was assessed in only a small number of courses. In contrast, analyzing student’s Wi-Fi digital traces made it possible to estimate class attendance at large-scale, i.e. across hundreds of courses with different start times. Additionally, we were able to control for unobserved student characteristics (e.g., effort and motivation) by comparing attendance rates for different start times in the same individuals. Within-student comparisons showed that attendance rates were higher for all other class start times relative to 08:00. We also provided the first objective evidence that students often sleep instead of going to early morning classes. Actigraphy data showed that students overslept for nearly one third of classes that started at 08:00. These results are consistent with survey studies of university students in which common reasons for absenteeism included lectures occurring too early, lack of sleep, feeling tired, and oversleeping (8-11).

Our findings suggest that morning classes contribute to a university-wide sleep debt. LMS data revealed differences in diurnal behavior for school days versus non-school days, which is consistent with previous studies that compared Twitter actions and smartphone interactions for weekdays and weekends (39-41). Students had shorter sleep opportunities on school days and the effects of class start times on LMS logins closely corresponded with actigraphy-derived sleep parameters. Our results showed that students went to bed at around the same time but woke up earlier to attend morning classes. Consequently, nocturnal sleep duration was shorter on nights that preceded earlier classes. Our results for objective sleep behavior extend prior work in undergraduates demonstrating that self-reported sleep duration was shorter when students’ first class of the day was earlier in the morning (27, 42).

Universities should avoid scheduling early morning classes if the goal is to minimize sleep deprivation and circadian misalignment. In our study, the midpoint of sleep occurred earlier for morning classes, indicating greater social jet lag. Sleep behavior on school days was similar to non-school days only when the first class of the day took place in the afternoon. Similar to our results, a prior study found that undergraduates with classes only in the morning (07:00-12:00) had shorter sleep on weekdays and extended their sleep duration on weekends, whereas students with classes only in the afternoon (14:00-18:00) did not show a difference in sleep duration between weekdays and weekends (43). Comparable results were obtained in adolescents attending schools with morning and afternoon/evening shifts, in which students with later class schedules obtained more sleep on school days and were less likely to extend their sleep on weekends (44-50). These findings for sleep behavior are consistent with models that predict much later start times are optimal for cognitive performance in undergraduates (51). Based on ratings of when students feel their best, the plateau of peak performance begins at around 11:00 or noon and extends into the late evening. A survey of student preferences at a pair of universities also found that the mean preferred start time was close to 11:00 (52). These studies suggest that early classes contribute to short sleep and circadian misalignment and should be discouraged.

We showed that having morning classes on more days of the week was associated with poorer academic performance. Previous studies found that grades were either lower in the morning compared with the afternoon (28-32), or there were no differences across time of day (53). These differences between studies could be related to the way that students were graded across courses and universities. At the university in our study, course grades are often adjusted to meet a recommended grade distribution (i.e., graded on a curve). By design, the average grade does not differ much between courses and may mask underlying performance differences between morning and afternoon courses. We therefore analyzed grade point average by considering students’ overall academic performance in relation to their frequency of morning classes. If morning classes have a cumulative negative impact on students’ attendance and sleep, performance should be lower for classes at other times of day. Consistent with this expectation, grade point average was highest for students with no morning classes and decreased when students had more days of morning classes. Similar to this result, a prior study found that students who had a first period course before 08:00 performed worse in all other courses taken on the same day, as compared with students who did not have a first period course (32). Notably, we found that the effect size of morning classes on grade point average was comparable in magnitude to many student- and teacher-related interventions for improving academic achievement (54). Hence, reducing the number of days that university students have morning classes could lead to meaningful improvements in learning and grade point average.

In our study, there are some limitations associated with using students’ digital traces to measure their behavior. While Wi-Fi connection data made it possible to measure attendance rates in large numbers of students and courses, this method requires that students use a Wi-Fi enabled device that is actively scanning for wireless access points. It is possible that some students disabled Wi-Fi scanning on their devices or used a cellular data plan instead. Nonetheless, we found that Wi-Fi confirmed attendance underestimated instructor-reported attendance by only a small amount, suggesting that Wi-Fi connection data provide a reliable relative estimate of the number of students who attend class. A limitation of using LMS data for estimating sleep behavior is that students typically interact with the LMS only a few times each day. Hence, the LMS method we used cannot be used to estimate daily sleep behavior in individual students. LMS usage patterns also differ between courses depending on the type of course and instructor preferences, and some students rarely use the LMS. Despite these limitations, we showed that LMS logins accumulate in large numbers over time and can provide universities with an aggregate view of how their class scheduling practices influence students’ sleep opportunities.

In conclusion, our study suggests that universities should avoid scheduling early morning classes to optimize students’ learning. While morning classes may be scheduled to maximize use of resources (classroom space and faculty) and to minimize scheduling conflicts for students and faculty, our results indicate that there is a trade-off whereby students are more likely to miss class, get less sleep, and obtain a lower grade point average. Early classes could be scheduled later in the day if classrooms and lecture theatres are not being fully utilized, and making classrooms a shared resource across departments might open up time slots for more afternoon/evening courses to be conducted in parallel. To justify taking such actions, universities need scalable methods for assessing the impact of their class scheduling practices on students. Our study showed that archived digital traces that are routinely collected by universities can be used to measure the impact of class start times on students’ behavior. In future studies, these approaches can be used to test the effectiveness of interventions for improving students’ class attendance, sleep, and academic achievement.

## Materials and Methods

### Ethics Statement

Permission to analyze university-archived data was obtained from the NUS Institute for Applied Learning Sciences and Educational Technology (ALSET). ALSET stores and links de-identified student data for educational analytics research on the ALSET Data Lake. University-archived datasets included students’ demographic information (age, sex, ethnicity, and year of matriculation), course enrolment, Wi-Fi connection data, use of the LMS, and grades. Analyses of these data were approved by the NUS Learning Analytics Committee on Ethics (LACE). Students whose university-archived data were included in our study provided informed consent to the NUS Student Data Protection Policy, which explains that their data can be used for research. Analyses of university-archived data were exempt from review by the NUS Institutional Review Board (IRB) because they were performed retrospectively on data that were de-identified to the researchers. Permission for collecting attendance data from course instructors was approved by LACE. Research procedures in the actigraphy study were approved by the NUS IRB and participants provided written informed consent to take part in the research and to add their data to the ALSET Data Lake.

### Wi-Fi confirmed attendance

Students’ Wi-Fi connection metadata were added to the ALSET Data Lake by NUS Information Technology. Each time that a student’s Wi-Fi enabled device associated with the NUS wireless network, the transmission data were logged. Data included the tokenized student identity, the anonymized media access control (MAC) address used to identify the Wi-Fi enabled device, the name and location descriptor of the Wi-Fi access point, and the start and end time of each Wi-Fi connection. The campus wireless network at NUS comprises more than 6,500 Wi-Fi access points, including coverage of classrooms and lecture halls (34). Students’ Wi-Fi connections at these locations were cross-referenced with their course timetables. These time and location data made it possible to identify students who connected to a Wi-Fi router in their classroom during class hours, thereby confirming their attendance.

The method of using Wi-Fi connection data to estimate class attendance was validated by collecting attendance data from course instructors. Attendance data were obtained for 53 class sessions across 13 different courses. In each of these class sessions, we determined the number of enrolled students with at least one Wi-Fi connection during class. The strength of the linear correlation between instructor-reported attendance and Wi-Fi confirmed attendance was assessed using Pearson’s correlation analysis.

Wi-Fi confirmed attendance was investigated over 3 semesters (2018/19 semester 1, 2018/19 semester 2, and 2019/20 semester 1) using all available data on the ALSET Data Lake prior to the COVID-19 pandemic. Courses were considered for the analysis if (1) they were categorized as a lecture course according to the NUS timetable, (2) they were held once per week, (3) they were held at least 7 times over the 13-week semester, (4) they lasted 2 h per session, and (5) they had at least 100 students enrolled in the course. The rationale for these inclusion criteria was to ensure that comparable types of courses were included in the analyses across different class start times. Among the 436 courses that met these criteria, 71 were excluded due to missing or incomplete Wi-Fi connection data or inconsistencies with the class timetable (e.g., due to cancelled or rescheduled classes). The remaining 365 courses were sorted by their start time, and data were analyzed only for those start times in which there were at least 5 courses per semester (08:00, 21 courses; 09:00, 18 courses; 10:00, 89 courses; 12:00, 67 courses; 14:00, 72 classes; 16:00, 70 classes). The final dataset included 337 courses and 24,678 unique students. The average class size (i.e., number of students enrolled in the course) in the dataset was 193 ± 73 students (mean ± SD), and class size did not differ between lecture start times (one-way ANOVA: *F*_5,331_ = 0.91, *P* = 0.476).

The Wi-Fi confirmed attendance rate for each student was determined in each of the 337 courses. In a given course, this was calculated as the number of lectures in which a student was detected by Wi-Fi, divided by the total number of lectures held during the semester. Due to heterogeneity in students’ course timetables, the number and combination of unique class start times differed between students (i.e., individual students did not have data for all class start times). We therefore used linear mixed-effects ANOVA to test the association between class start time and Wi-Fi confirmed attendance. Class start time was entered as a repeated fixed factor with student included as a random factor. In instances where students took more than one course with the same start time (on different days), the median Wi-Fi confirmed attendance rate across courses was entered into the model (i.e., each student contributed one data point for a given start time). The linear mixed-effects model was implemented using the “lmerTest” package (version 3.1-3) with R Statistical Software (version 3.6.3) (55). An F-test was used to test for fixed effects (threshold for statistical significance, *P* < 0.05), with degrees of freedom estimated using Satterthwaite’s method. Multiple comparisons between the 6 different class start times were performed using paired student t-tests with a Bonferroni correction (15 pairwise comparisons, P < 0.0033). Effect sizes and permutated p-values were calculated with the “dabest” package (version 0.3.0) using Python 3.7.8 (56).

### Learning Management System (LMS) data

Students’ logins on the university’s LMS were analyzed over 5 semesters (2016/17 semester 2, 2017/18 semester 1, 2017/18 semester 2, 2018/19 semester 1, 2018/19 semester 2), using all available data on the ALSET Data Lake. The diurnal time courses of logins were analyzed separately in each semester by sorting the data according to each student’s first class start time of the day. Analyses were restricted to the most frequent class start times at NUS for which we also had sufficient actigraphy data for making comparisons (08:00, 09:00, 10:00, 11:00, 12:00, 14:00 and 16:00). Data were also analyzed on non-school days, corresponding to weekends and weekdays with no scheduled classes. For a given class start time, the total number of logins per 5-min bin were summed across all students starting from 19:00 on the previous evening until 19:00 in the evening of the day in which the class took place (288 epochs per day). The same 24-h time window was examined on non-school days. The aggregated time series data allowed us to compare students’ diurnal login behavior by their first class of the day, and relative to days with no classes. The dataset comprised 17.4 million logins from 39,458 students.

The time courses of logins in each semester were used to derive the offset and onset of LMS activity. These parameters were determined using a threshold crossing method. Each time series was normalized by dividing the number of logins in each 5-min bin by the total number of logins across all bins (i.e., the sum across all bins was set to a value of 1). The LMS activity threshold was calculated as 50% of the average normalized number of logins per bin (normalized threshold = 1 bin / 288 bins × 0.5 = 0.001736). The LMS login offset was defined as the clock time when the normalized number of logins dropped and remained below this threshold. The LMS login onset was defined as the time point when logins exceeded and stayed above threshold. The LMS inactive period was defined as the duration of time from the LMS login offset to the LMS login onset. The midpoint of the LMS inactive period was also calculated between the LMS login offset and onset. Separate ANOVAs with the repeated factor semester (5 different semesters) were used to test the association of school start time (08:00, 09:00, 10:00, 11:00, 12:00, 14:00, 16:00, No class) with each LMS-derived parameter (offset, onset, inactive period, midpoint). Multiple comparisons of LMS activity between each class start time and days with no classes were performed using paired t-tests with a Bonferroni correction (7 pairwise comparisons, *P* < 0.0071). ANOVA was performed using R Statistical Software (version 3.6.3).

### Actigraphy study

NUS undergraduates aged 18-25 years were recruited to take part in a 6-week research study of their sleep-wake patterns during the school term. Participants were required to be non-smokers in good general health with a body mass index between 18.5-27.0 kg/m^2^. Individuals were ineligible if they reported shift work (paid work between 23:00 and 07:00) or if they planned on traveling across time zones during the study. Participants wore an actigraphy watch (Actiwatch Spectrum Plus or Actiwatch 2; Philips Respironics Inc., Pittsburgh, PA) on their non-dominant hand and made weekly visits to a classroom to have their data downloaded and to undergo a set of neurobehavioral tests (results not reported here). Among 202 undergraduate students who enrolled in the study, 13 participants withdrew before the end of the data collection period (no longer available, n = 6; personal reasons, n = 5; falling ill, n = 2), and 5 participants were withdrawn from the study by the researchers for not complying with study procedures (e.g., not wearing the actigraphy watch or not showing up on time for appointments). In the remaining 184 participants who wore the actigraphy watch for 6 weeks, 2 individuals were excluded because of poor quality data, and 1 individual failed to provide his course timetable with his class start times. The final dataset included 181 student participants with 7,312 nocturnal sleep recordings (range, 27-42 days per individual).

Students were instructed to wear the actigraphy watch at all times except when taking part in activities that might damage the device (e.g., contact sports or swimming). Participants pressed an event marker button on their watch when going to bed/ waking up and when putting on/ taking off the actigraphy watch. They were also required to complete a daily diary of times that they slept or removed the actigraphy watch. Actigraphy data were collected in 30-s epochs and analyzed using Actiware software (version 6.0.9). Time-in-bed intervals were marked in the actogram using participants’ event marker presses and sleep diary entries. Event markers were prioritized over diary entries in instances where there was poor agreement between measures, while also taking into account each individual’s pattern of activity and light exposure. Actograms were inspected, reviewed, and approved by all members of the research team before analyzing the data to derive sleep variables. Sleep scoring of each time-in-bed interval was performed using the medium wake-sensitivity threshold (threshold = 40 activity counts) and a 10-min immobility threshold for determining sleep onset and sleep offset.

The primary sleep variables were (1) sleep onset, (2) sleep offset, (3) nocturnal total sleep time, (4) nocturnal time in bed for sleep, and (5) midpoint of the sleep period. Each student’s actigraphy data were sorted by his/her first class of the day, and we restricted our analyses to class times in which there were at least 20 individuals whose first class of the day started at that time (08:00, n = 103; 09:00, n = 61; 10:00, n = 123; 11:00, n = 35; 12:00, n = 107; 14:00, n = 71; 16:00, n = 44). The frequency of instances in which students failed to wake up in time for class was evaluated by pooling data across participants for a given class start time. Chi-squared tests were used to test for differences across class start times in the frequency of (1) waking up after the start of class and (2) waking up too late to reach class on time, which took into account travel time to reach school. The latter was assessed using the question “How long does it usually take for you to get from your residence to your first class of the day?”. The dataset comprised 6,546 sleep offset values from 174 participants who reported their travel time (Start time of first class, number of sleep offset values: 08:00, 776 values; 09:00, 468 values; 10:00, 940 values; 11:00, 169 values; 12:00, 631 values; 14:00, 389 values; 16:00, 164 values; No class, 3,009 values).

Linear mixed-effects ANOVA was used to test associations of students’ first class time of the day (08:00, 09:00, 10:00, 11:00, 12:00, 14:00, 16:00, No class) with each actigraphy-derived sleep variable. After sorting each student’s actigraphy data by his/her first class of the day, the median value for each sleep variable was determined for a given class start time. For example, if a student had classes on 3 days per week starting at 08:00, 10:00, and 12:00, the median values for sleep onset, sleep offset, total sleep time, time in bed, and midpoint of sleep were calculated separately for each of the respective start times over the 6-week recording interval. Class start time was entered in the model as a repeated fixed factor with student included as a random factor. The linear mixed-effects model was implemented using the “lmerTest” package (version 3.1-3) in R Statistical Software (version 3.6.3) (55). Multiple comparisons for sleep variables between each class start time and days with no class were performed using paired t-tests with a Bonferroni correction (7 pairwise comparisons, *P* < 0.0071). Effect sizes and permutated p-values were calculated with the “dabest” package (version 0.3.0) using Python 3.7.8 (56).

### Academic performance

Students’ course grades were analyzed over the 6 semesters that Wi-Fi connection data and LMS data were available (2016/17 semester 2, 2017/18 semester 1, 2017/18 semester 2, 2018/19 semester 1, 2018/19 semester 2, 2019/20 semester 1). At NUS, students are given a letter grade that is converted to a number for calculating the grade point (A+ = 5.0, A = 5.0, A- = 4.5, B+ = 4.0, B = 3.5, B- = 3.0, C+ = 2.5, C = 2.0, D+ = 1.5, D = 1.0, F = 0.0). Students earn course credits based on the estimated workload hours per week, and the grade point average represents the cumulative performance weighted by the credits earned in each course. Because a course can have multiple class start times (e.g., 10:00 lecture on Monday, and 16:00 tutorial on Wednesday), we decided to group data by morning and afternoon courses. Morning courses were defined as having all classes (e.g., lectures, tutorials, and laboratories) start before 12:00, and afternoon courses were defined as having all classes start at 12:00 or later. Courses that had class meetings in the morning and afternoon were excluded from the analysis. In each semester, we restricted our analyses to students who earned 20 course credits (the mode of the distribution for course credits) to ensure that they had a comparable total workload. This usually corresponded to taking 4 or 5 courses concurrently. The final sample included 27,281 unique students, ranging from 9,220 to 11,854 students per semester. The dataset comprised 129,919 individual course grades.

The distribution of grades for morning and evening courses was compared (in each semester and using aggregated data) using a Mann-Whitney U test. The effect size of taking courses in the morning versus the afternoon on grades was measured using Cliff’s delta because grade point data were ordinal. In separate analyses, we calculated the grade point average for students grouped by the number of days per week that they had a morning class (0, 1, 2, 3, 4, or 5 days). The grade point average was calculated using all grades that a student obtained during a given semester, irrespective of the timing that classes occurred. Analyses were restricted to students taking 20 course credits. ANOVA was used to test the association of days per week with morning classes with grade point average. Effect size was measured using Cohen’s d, with the reference category defined as students with no morning classes. Effect sizes and permutated p-values were calculated with the “dabest” package (version 0.3.0) using Python 3.7.8 (56).

## Acknowledgments and funding sources

We thank researchers and students in the Chronobiology and Sleep Laboratory for collecting data; researchers and administrative staff at the NUS Institute for Applied Learning Sciences and Educational Technology (ALSET) and NUS Information Technology (NUS IT) for supporting analyses of university-archived data; and Fun Man Fung and Edwin Setiadi Sugeng for facilitating the collection of attendance data. Data storage and management were supported by the NUS Office of the Senior Deputy President & Provost and ALSET. The work was funded by the Ministry of Education, Singapore (MOE2019-T2-2-074) and the National Research Foundation, Singapore (NRF2016-SOL002-001).

## Supplementary Information for

**Fig. S1.**
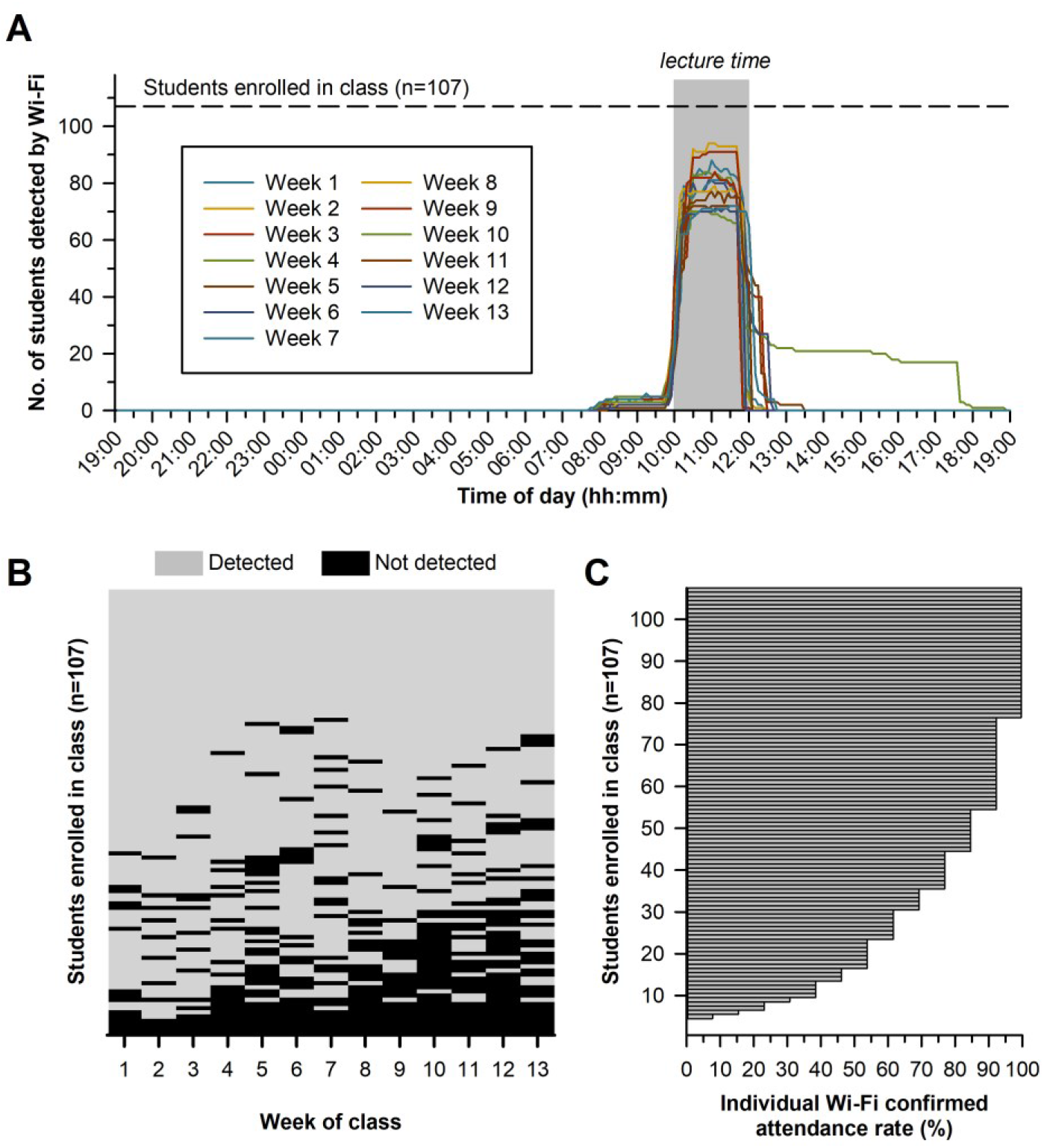
Method for using Wi-Fi connection data to estimate students’ class attendance. Wi-Fi confirmed attendance data are shown for a representative lecture course with a class enrolment of 107 students. The class took place weekly from 10:00 to 12:00 over 13 weeks. (**A**) Daily time courses are shown for the number of enrolled students who connected to a Wi-Fi access point in the lecture hall where the class was conducted. The number of students detected by Wi-Fi increased sharply at the start of each class and then dropped at the end of class. There was one week when a group of about 20 students remained in the lecture hall for several hours after class. (**B**) The binary heat map shows individually-determined Wi-Fi confirmed attendance (detected or not detected during class hours) across all 13 weeks of the semester. (**C**) Wi-Fi confirmed attendance rates are shown for individual students in the course. Each row in panels B and C represents the same individuals.

**Fig. S2.**
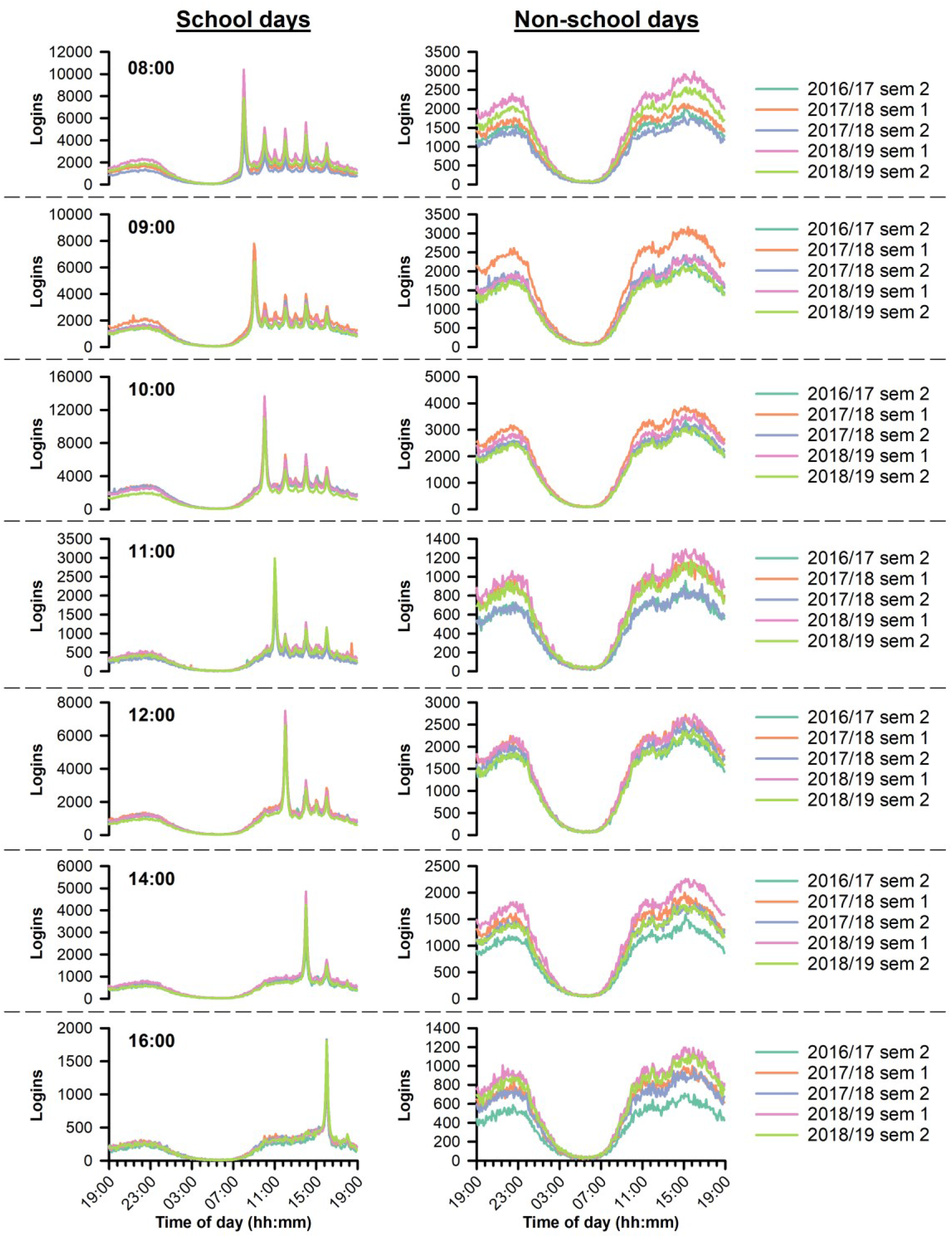
Daily time courses of Learning Management System (LMS) logins. Students’ LMS logins were compiled by time of day in 5-min bins over 5 different semesters. Data were sorted by students’ first class of the day (left column) and compared with non-school days (right column) in the same group of students. The time courses of LMS logins were highly reproducible across semesters and academic years.

**Fig. S3.**
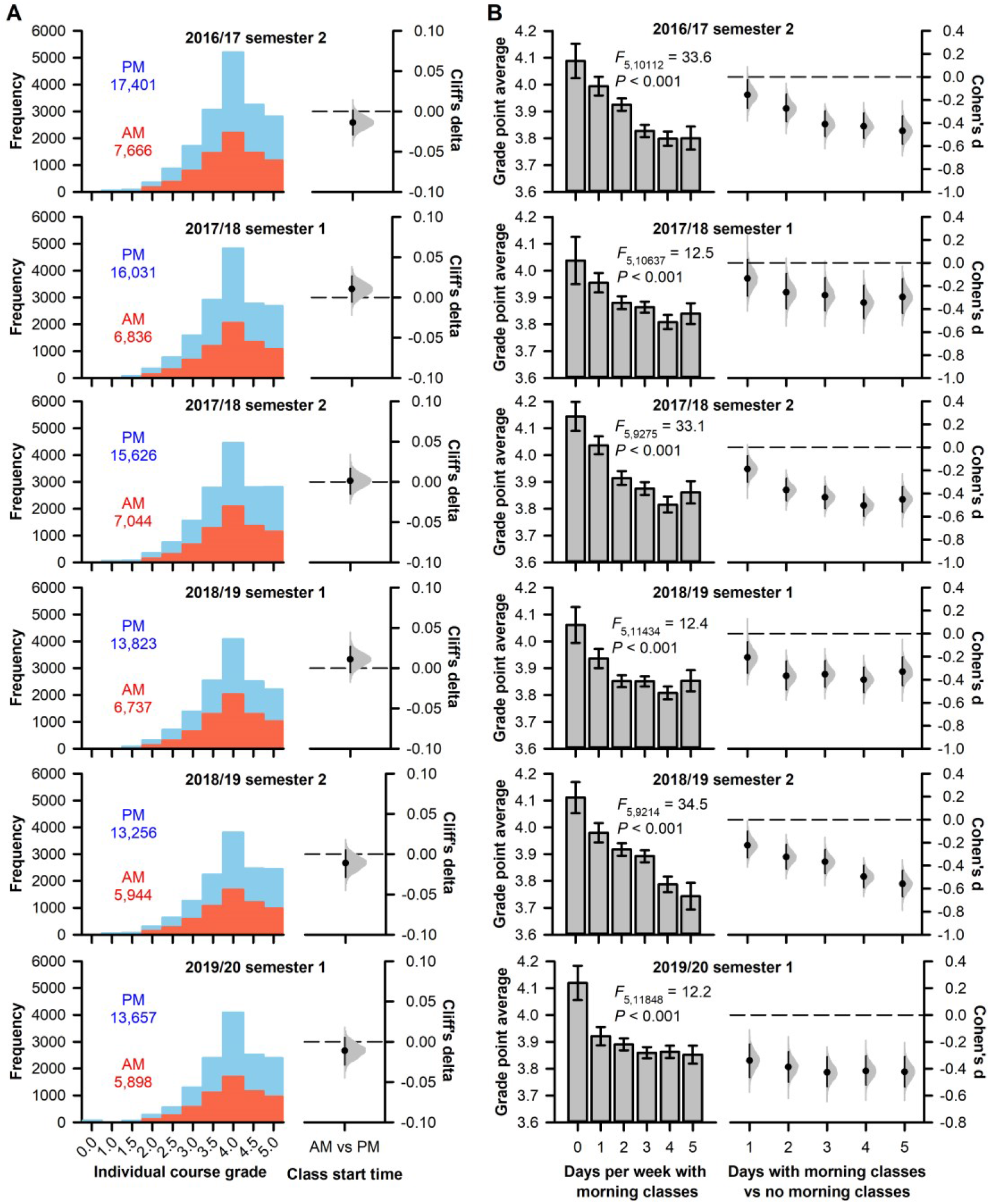
Effects of morning classes on students’ academic performance. (**A**) The distributions of grades are shown for courses that took place exclusively in the morning (starting before 12:00) or exclusively in the afternoon (starting at 12:00 or later) over 6 semesters. The number of morning and afternoon course grades is indicated on each plot. Effect sizes (Cliff’s delta) of morning courses on grades are shown relative to afternoon courses. (**B**) In all 6 semesters, grade point average was lower in students who had morning classes on more days of the week. The mean ± 95% CIs are shown for students categorized by the number of days that they had morning classes. Effect sizes (Cohen’s d) with 95% CIs and the bootstrap sampling distributions are plotted for days with morning classes (1 to 5 days) versus having no morning classes.

**Table S1.** Demographic characteristics of students included in different analyses.

|  | 2016/17<br>semester 2 | 2017/18<br>semester 1 | 2017/18<br>semester 2 | 2018/19<br>semester 1 | 2018/19<br>semester 2 | 2019/20<br>semester 1 |
| --- | --- | --- | --- | --- | --- | --- |
| **Wi-Fi connection data (lectures with ≥100 students enrolled)** |  |  |  |  |  |  |
| n | - | - | - | 18,691 | 17,951 | 20,554 |
| Age (mean ± SD) | - | - | - | 21.1 ± 1.9 | 21.5 ± 1.9 | 21.3 ± 2.0 |
| Female, n (%) | - | - | - | 9,345 (50.0%) | 9,035 (50.3%) | 10,736 (52.2%) |
| Chinese, n (%) | - | - | - | 16,231 (86.8%) | 15,601 (86.9%) | 17,779 (86.5%) |
| Class year, n (%) |  |  |  |  |  |  |
| 1 | - | - | - | 6,346 (34.0%) | 6,356 (35.4%) | 5,099 (24.8%) |
| 2 | - | - | - | 5,590 (29.9%) | 5,330 (29.7%) | 6,333 (30.8%) |
| 3 | - | - | - | 3,871 (20.7%) | 3,644 (20.3%) | 4,547 (22.1%) |
| 4 | - | - | - | 2,611 (14.0%) | 2,502 (13.9%) | 4,105 (20.0%) |
| 5+ | - | - | - | 273 (1.5%) | 119 (0.7%) | 470 (2.3%) |
| **Learning management system data** |  |  |  |  |  |  |
| n | 22,580 | 24,615 | 22,996 | 25,506 | 24,121 | - |
| Age (mean ± SD) | 21.8 ± 2.3 | 21.5 ± 2.5 | 21.9 ± 2.4 | 21.5 ± 2.6 | 21.9 ± 2.5 | - |
| Female, n (%) | 11,829 (52.4%) | 12,586 (51.1%) | 11,823 (51.4%) | 12,979 (50.9%) | 12,343 (51.2%) | - |
| Chinese, n (%) | 19,661 (87.1%) | 21,305 (86.6%) | 19,934 (86.7%) | 22,019 (86.3%) | 20,840 (86.4%) | - |
| Class year, n (%) |  |  |  |  |  |  |
| 1 | 6,417 (28.4%) | 67,41 (27.4%) | 6,577 (28.6%) | 7,404 (29.0%) | 7,365 (30.5%) | - |
| 2 | 6,122 (27.1%) | 6,420 (26.1%) | 5,980 (26.0%) | 6,625 (26.0%) | 6,268 (26.0%) | - |
| 3 | 5,027 (22.3%) | 5,170 (21.0%) | 4,984 (21.7%) | 5,154 (20.2%) | 4,913 (20.4%) | - |
| 4 | 4,611 (20.4%) | 5,532 (22.5%) | 5,073 (22.1%) | 5,508 (21.6%) | 5,198 (21.5%) | - |
| 5+ | 403 (1.8%) | 752 (3.1%) | 382 (1.7%) | 815 (3.2%) | 377 (1.6%) | - |
| **Grades data (students with 20 course credits)** |  |  |  |  |  |  |
| n | 10,118 | 10,643 | 9,281 | 11,440 | 9,220 | 11,854 |
| Age (mean ± SD) | 21.3 ± 1.8 | 20.9 ± 1.8 | 21.4 ± 1.7 | 20.8 ± 1.8 | 21.3 ± 1.8 | 20.7 ± 1.8 |
| Female, n (%) | 5,923 (58.5%) | 5,850 (55.0%) | 5,184 (55.9%) | 6,315 (55.2%) | 5,016 (54.4%) | 6,132 (51.7%) |
| Chinese, n (%) | 8,814 (87.1%) | 9,343 (87.8%) | 8,065 (86.9%) | 9,942 (86.9%) | 7,940 (86.1%) | 10,102 (85.2%) |
| Class year, n (%) |  |  |  |  |  |  |
| 1 | 3,125 (30.9%) | 4,163 (39.1%) | 2,566 (27.6%) | 4,964 (43.4%) | 2,992 (32.5%) | 5,252 (44.3%) |
| 2 | 3,793 (37.5%) | 2,954 (27.8%) | 3,527 (38.0%) | 3,234 (28.3%) | 3,451 (37.4%) | 3,647 (30.8%) |
| 3 | 1,964 (19.4%) | 2,241 (21.1%) | 1,768 (19.0%) | 2,073 (18.1%) | 1,558 (16.9%) | 1,848 (15.6%) |
| 4 | 1,177 (11.6%) | 1,132 (10.6%) | 1,366 (14.7%) | 1,025 (9.0%) | 1,155 (12.5%) | 900 (7.6%) |
| 5+ | 59 (0.6%) | 153 (1.4%) | 54 (0.6%) | 144 (1.3%) | 64 (0.7%) | 207 (1.7%) |

**Table S2.**
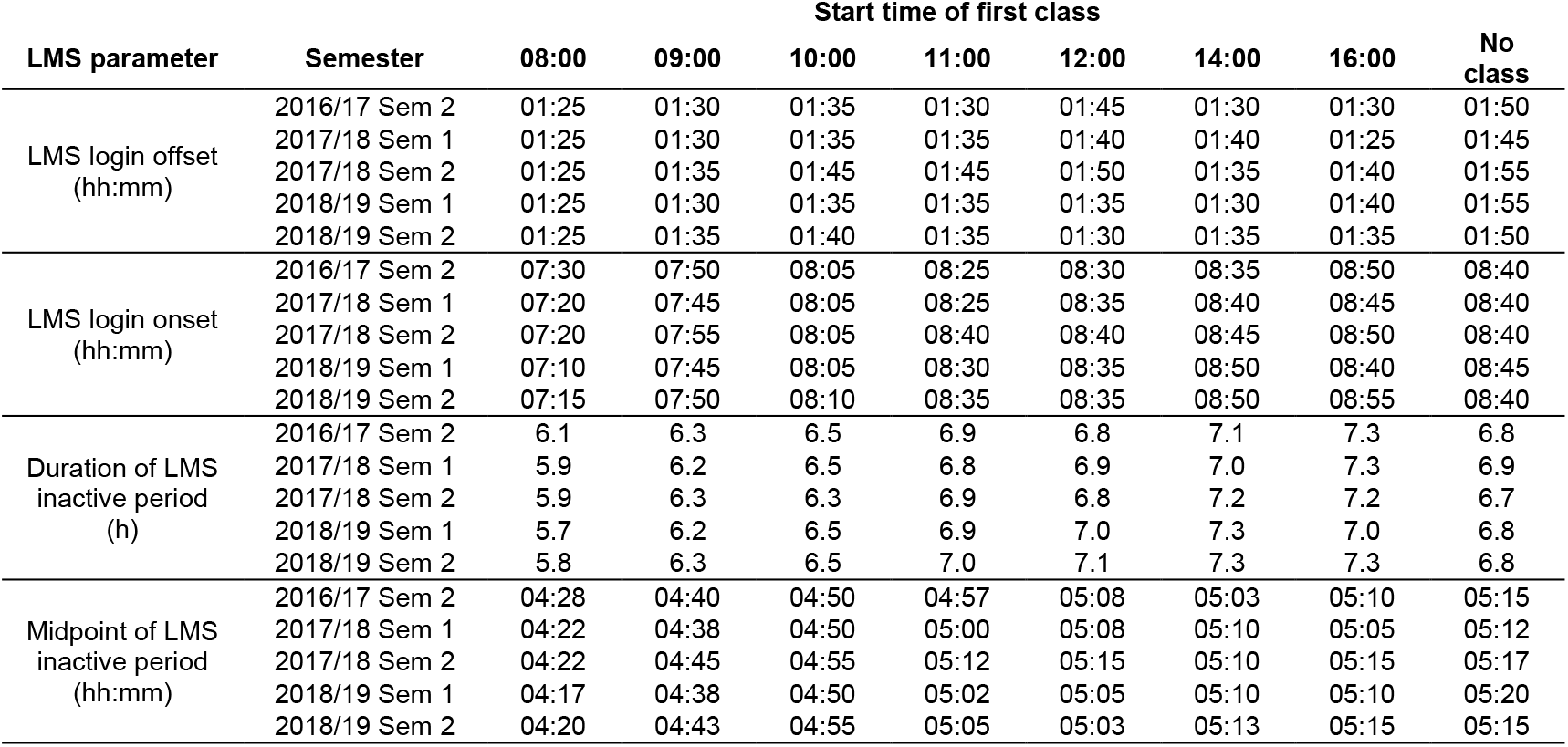
Learning Management System (LMS)-derived parameters sorted by students’ first class of the day.

**Table S3.** Actigraphy-derived sleep parameters sorted by students’ first class of the day. The mean ± SD is shown for each sleep variable. TST = total sleep time; TIB = Time in bed.

| Start time of first class (hh:mm) | n | Sleep onset (hh:mm) | Sleep offset (hh:mm) | Nocturnal TST (h) | Nocturnal TIB (h) | Daily TST (h) | Daily TIB (h) |
| --- | --- | --- | --- | --- | --- | --- | --- |
| 08:00 | 103 | 01:35 ± 01:13 | 07:18 ± 01:18 | 5.2 ± 1.1 | 6.2 ± 1.3 | 5.3 ± 1.1 | 6.4 ± 1.3 |
| 09:00 | 61 | 01:50 ± 01:14 | 07:56 ± 01:06 | 5.5 ± 1.0 | 6.7 ± 1.1 | 5.6 ± 1.0 | 6.8 ± 1.1 |
| 10:00 | 123 | 01:58 ± 01:20 | 08:11 ± 00:58 | 5.6 ± 0.9 | 6.8 ± 1.0 | 5.7 ± 1.0 | 6.9 ± 1.0 |
| 11:00 | 35 | 02:21 ± 01:26 | 08:37 ± 01:04 | 5.7 ± 1.3 | 6.9 ± 1.3 | 5.8 ± 1.2 | 7.1 ± 1.1 |
| 12:00 | 107 | 01:48 ± 01:18 | 08:37 ± 01:13 | 6.2 ± 0.9 | 7.4 ± 1.1 | 6.2 ± 0.9 | 7.5 ± 1.1 |
| 14:00 | 71 | 02:02 ± 01:36 | 08:49 ± 01:26 | 6.0 ± 1.2 | 7.4 ± 1.4 | 6.1 ± 1.1 | 7.4 ± 1.4 |
| 16:00 | 44 | 01:33 ± 01:23 | 08:54 ± 01:21 | 6.4 ± 1.0 | 7.8 ± 1.3 | 6.5 ± 1.1 | 7.8 ± 1.3 |
| No class | 181 | 01:59 ± 01:14 | 09:04 ± 01:18 | 6.3 ± 0.9 | 7.6 ± 1.0 | 6.4 ± 0.9 | 7.7 ± 1.0 |

**Table S4.** Effect sizes of different class start times on sleep relative to days with no classes.

| Sleep variable | Start time of first class (hh:mm) | Change relative to non-school days |  |
| --- | --- | --- | --- |
|  |  | Mean difference in hours (95% CI) | Cohen's d (95% CI) |
| Sleep onset | 08:00 | -0.38 (-0.55 to -0.22) | -0.30 (-0.44 to -0.17) |
|  | 09:00 | -0.27 (-0.48 to -0.05) | -0.21 (-0.39 to -0.03) |
|  | 10:00 | -0.11 (-0.27 to 0.05) | -0.08 (-0.21 to 0.04) |
|  | 11:00 | -0.21 (-0.46 to 0.01) | -0.15 (-0.37 to 0.01) |
|  | 12:00 | -0.16 (-0.30 to -0.02) | -0.13 (-0.24 to -0.01) |
|  | 14:00 | 0.05 (-0.16 to 0.26) | 0.03 (-0.12 to 0.18) |
|  | 16:00 | -0.29 (-0.50 to -0.06) | -0.21 (-0.40 to -0.04) |
| Sleep offset | 08:00 | -1.72 (-1.99 to -1.43) | -1.29 (-1.59 to -1.00) |
|  | 09:00 | -1.12 (-1.44 to -0.81) | -0.90 (-1.23 to -0.53) |
|  | 10:00 | -0.94 (-1.17 to -0.75) | -0.85 (-1.07 to -0.62) |
|  | 11:00 | -0.97 (-1.37 to -0.60) | -0.75 (-1.08 to -0.46) |
|  | 12:00 | -0.55 (-0.72 to -0.38) | -0.43 (-0.56 to -0.30) |
|  | 14:00 | -0.20 (-0.42 to 0.00) | -0.14 (-0.31 to 0.01) |
|  | 16:00 | -0.25 (-0.49 to -0.03) | -0.19 (-0.38 to -0.02) |
| Nocturnal time in bed for sleep | 08:00 | -1.35 (-1.57 to -1.12) | -1.14 (-1.39 to -0.89) |
|  | 09:00 | -0.80 (-1.08 to -0.53) | -0.74 (-1.04 to -0.45) |
|  | 10:00 | -0.83 (-1.02 to -0.64) | -0.80 (-1.00 to -0.60) |
|  | 11:00 | -0.56 (-0.95 to -0.08) | -0.50 (-0.85 to -0.02) |
|  | 12:00 | -0.43 (-0.64 to -0.24) | -0.42 (-0.61 to -0.22) |
|  | 14:00 | -0.24 (-0.48 to 0.00) | -0.19 (-0.39 to 0.00) |
|  | 16:00 | -0.16 (-0.45 to 0.11) | -0.14 (-0.39 to 0.11) |
| Nocturnal total sleep time | 08:00 | -1.16 (-1.37 to -0.95) | -1.13 (-1.37 to -0.89) |
|  | 09:00 | -0.70 (-0.94 to -0.47) | -0.74 (-1.02 to -0.44) |
|  | 10:00 | -0.70 (-0.88 to -0.53) | -0.74 (-0.93 to -0.55) |
|  | 11:00 | -0.53 (-0.89 to -0.15) | -0.48 (-0.78 to -0.11) |
|  | 12:00 | -0.31 (-0.49 to -0.13) | -0.34 (-0.54 to -0.14) |
|  | 14:00 | -0.17 (-0.37 to 0.02) | -0.16 (-0.36 to 0.02) |
|  | 16:00 | -0.12 (-0.37 to 0.13) | -0.13 (-0.39 to 0.14) |
| Midpoint of sleep | 08:00 | -1.02 (-1.21 to -0.83) | -0.89 (-1.12 to -0.67) |
|  | 09:00 | -0.59 (-0.82 to -0.36) | -0.51 (-0.74 to -0.25) |
|  | 10:00 | -0.47 (-0.62 to -0.33) | -0.43 (-0.60 to -0.27) |
|  | 11:00 | -0.47 (-0.68 to -0.24) | -0.40 (-0.62 to -0.20) |
|  | 12:00 | -0.35 (-0.46 to -0.23) | -0.30 (-0.40 to -0.20) |
|  | 14:00 | -0.08 (-0.25 to 0.09) | -0.07 (-0.20 to 0.07) |
|  | 16:00 | -0.28 (-0.44 to -0.10) | -0.23 (-0.39 to -0.08) |

